# SEREEGA: Simulating Event-Related EEG Activity

**DOI:** 10.1101/326066

**Authors:** Laurens R. Krol, Juliane Pawlitzki, Fabien Lotte, Klaus Gramann, Thorsten O. Zander

**Affiliations:** Biological Psychology and Neuroergonomics, Technische Universität Berlin, Sekr. KWT-1, Fasanenstr. 1, 10623 Berlin, Germany; Zander Laboratories B.V., Amsterdam, The Netherlands; Inria, LaBRI (CNRS/Univ Bordeaux/Bordeaux INP), Talence, France

**Keywords:** Electroencephalography, Simulation, Methodology, Evaluation, Ground Truth

## Abstract

Electroencephalography (EEG) is a popular method to monitor brain activity, but it can be difficult to evaluate EEG-based analysis methods because no ground-truth brain activity is available for comparison. Therefore, in order to test and evaluate such methods, researchers often use simulated EEG data instead of actual EEG recordings, ensuring that it is known beforehand which e ects are present in the data. As such, simulated data can be used, among other things, to assess or compare signal processing and machine learning algorithms, to model EEG variabilities, and to design source reconstruction methods. In this paper, we present SEREEGA, short for *Simulating Event-Related EEG Activity*. SEREEGA is a MATLAB-based open-source toolbox dedicated to the generation of simulated epochs of EEG data. It is modular and extensible, at initial release supporting five different publicly available head models and capable of simulating multiple different types of signals mimicking brain activity. This paper presents the architecture and general work ow of this toolbox, as well as a simulated data set demonstrating some of its functions.

**Highlights:**

- Simulated EEG data has a known ground truth, which can be used to validate methods.
- We present a general-purpose open-source toolbox to simulate EEG data.
- It provides a single framework to simulate many different types of EEG recordings.
- It is modular, extensible, and already includes a number of head models and signals.
- It supports noise, oscillations, event-related potentials, connectivity, and more.

## 1 Introduction

Having seen almost a century of continuous research and development since its first application on humans in the 1920s (Berger, 1929), electroencephalography (EEG) is now widely used in, among others, clinical settings, neuroscience, cognitive science, psychophysiology, and brain-computer interfacing, while its use continues to expand in fields such as neuroergonomics (Parasuraman & Rizzo, 2007; Frey, Daniel, Castet, Hachet, & Lotte, 2016), neurogaming (Krol, Freytag, & Zander, 2017), neuromarketing (Vecchiato et al., 2011), neuroadaptive technology (Zander, Krol, Birbaumer, & Gramann, 2016) and mobile brain/body imaging (Gramann et al., 2011). As of December 2017, PubMed reported over 140 000 publications related to EEG, with over 4 000 published in each of the past five years.

EEG reflects the electric fields that arise primarily due to the synchronous activity of post-synaptic potentials at apical dendrites in the cortical surface of the brain, as recorded by electrodes placed on the scalp. As such, it measures a specific subset of brain activity. This enables cognitive and affective correlates to be found in the EEG, allowing post-recording or even real-time evaluation of certain mental states exhibited by the recorded person, such as surprise (Donchin, 1981), error perception (Falkenstein, Hohnsbein, Hoormann, & Blanke, 1990; Blankertz, Schäfer, Dornhege, & Curio, 2002), task load (Klimesch, 1999; Mühl, Jeunet, & Lotte, 2014; Zander, Shetty, et al., 2017), or imagined movement (Pfurtscheller & Neuper, 2001; Blankertz, Dornhege, Krauledat, Müller, & Curio, 2007). Compared to other brain monitoring and imaging methods, EEG is relatively inexpensive and provides a high temporal resolution. It is also becoming increasingly portable and ready for personal use (Zander, Andreessen, et al., 2017) in various forms of neuroadaptive technology (Krol, Andreessen, & Zander, 2018). This explains EEG’s relative popularity.

However, these advantages present a trade-off, with costs incurred primarily in spatial resolution. A single electrode records the average activity of up to a billion neurons, and probably never less than 10 million neurons (Nunez & Srinivasan, 2006). This and other issues including volume conduction, the placement and distance of the electrodes relative to the cortical genserators of the activity they measure, and the complex relation between cortical functions and features of scalp potentials, require that great care is taken when analysing and interpreting raw EEG recordings.

A host of methods have been developed over the last decades to extract robust features from the recorded EEG that correlate to cortical functions and can be understood in a neurophysiological or statistical sense. Examples are the event-related potential technique (Luck, 2014), independent component analysis (Makeig, Jung, Bell, & Sejnowski, 1996), common spatial patterns (Guger, Ramoser, & Pfurtscheller, 2000), hierarchical linear modelling (Pernet, Chauveau, Gaspar, & Rousselet, 2011), and a many signal processing and machine learning algorithms (Lotte, Congedo, Lécuyer, Lamarche, & Arnaldi, 2007).

One major difficulty in developing techniques for EEG analyses is that there is no *ground truth* that describes the exact brain activity: The recorded EEG data cannot be compared to the actual neuro-electric activity of the brain because no technique exists to provide such a reference measurement. Thus, developers must use other ways to examine the validity of their EEG analysis approaches. The exact nature and detail of the required ground truth of course di ers depending on the analysis.

To that end, simulated EEG data (or “toy data”) has often been used to test and validate methods, as for example with blind source separation (Makeig, Jung, Ghahremani, & Sejnowski, 1999; Potter, Gadhok, & Kinsner, 2002), connectivity measures (Silfverhuth, Hintsala, Kortelainen, & Seppänen, 2012; Stam, Nolte, & Daffertshofer, 2007), artefact removal (Romo-Vazquez, Ranta, Louis-Dorr, & Maquin, 2007; He, Wilson, Russell, & Gerschutz, 2007), functional brain imaging (Gramfort, Strohmeier, Haueisen, Hamalainen, & Kowalski, 2011) and neurophysiological weight vector interpretation (Haufe et al., 2014). The authors of these examples all implemented simulation approaches from scratch, usually by linearly mixing a number of independent signals. This linear mixing was done using random weights, i.e., no realistic spatial information was taken into account. Other approaches use head models in order to provide more realistic linear mixing and add spatial dependencies to the simulation (e.g., Giraldo, den Dekker, & Castellanos-Dominguez, 2010; Haufe, Tomioka, Nolte, Müller, & Kawanabe, 2010; Haufe, Nikulin, Müller, & Nolte, 2013; Huiskamp, 2008).

Such custom-made approaches are often difficult to reproduce, as they have been implemented using different software packages, are reported at different levels of abstraction, and/or may be using head models that are not publicly available.

Some authors have not implemented their own methods, but have instead relied on commercially available packages (e.g., Yao & Dewald, 2005; Lansbergen, van Dongen-Boomsma, Buitelaar, & Slaats-Willemse, 2011). These packages, however, are not specifically designed for the purpose of simulation, nor are they freely available to the general public or open to scrutiny.

One open-source package known to the authors that provides simulation functionality is the Source Information Flow Toolbox (SIFT; Delorme et al., 2011). Its main purpose is to investigate structural, functional, and effective connectivity between brain regions and networks, but the toolbox can also be used to simulate scalp EEG using a system of coupled oscillators. Such simulations can serve as effective ground truth for the same connectivity measures that the toolbox investigates, and have been used to evaluate blind source separation methods as well (Hsu, Mullen, Jung, & Cauwenberghs, 2014). The SIFT toolbox, however, is not intended for wider-purpose simulation and as such, the methods that can be meaningfully applied to the data it can generate are limited.

In a more normative approach, Haufe and Ewald (2016) proposed a simulation and evaluation framework and made their code publicly available. The framework includes forward modelling using a realistic head model, both source and sensor noise with controlled signal-to-noise ratios, and signal generation based on autoregressive models. As with SIFT, this approach focuses solely on connectivity measures on continuous data, and thus provides no further signal generation options.

A final EEG simulation approach of note is the *phantom head* created by Oliveira, Schlink, Hairston, Knig, and Ferris (2016). They constructed a physical head model, filling a mannequin head with a conductive plaster and inserting antennae to simulate current dipoles. This allows unique effects to be investigated: For example, the authors investigated the effects of head motion by placing the model on a motion platform. Constructing such hardware, however, is not a viable approach for most questions where simulation can provide an answer. Importantly, software is easier to share, maintain, adapt, and extend.

Clearly, EEG data simulation is widely used as a tool to assess and validate the methods that are in use and in development. However, to the best of the authors’ knowledge, there currently exists no software package whose sole or primary purpose it is to simulate different types of EEG data, i.e. a dedicated, general-purpose EEG data simulation tool. We therefore present SEREEGA, short for *Simulating Event-Related EEG Activity*, making EEG data simulation more accessible to researchers.

SEREEGA is an open-source, MATLAB-based toolbox to generate simulated, event-related EEG data. Starting with a forward model obtained from a head model or pre-generated lead field, dipolar *brain components* can be defined. Each component has a specified position and orientation in the head model. Different activation patterns or *signals* can be assigned to these components. Scalp EEG data is simulated by projecting all signals from all components onto the scalp and summing these projections together.

SEREEGA is modular in that different head models and lead fields can be supported, as well as different activation signals. Five lead fields are currently supported, four pre-generated, the fifth customisable according to the user’s needs from a standard head model. Five types of activation signals are provided, allowing the simulation of different types of systematic (event-related) activity in both the time and the frequency domain, as well as the inclusion of any already existing time series as an activation signal.

This toolbox is intended to be a tool to generate data with a known ground truth in order to evaluate neuroscientific and signal processing methods, such as blind source separation, source localisation, connectivity measures, brain-computer interface classifier accuracy, derivative EEG measures, et cetera.

In the following, we first introduce the architecture of the toolbox and provide a brief introduction to its basic functionality and workflow. We then provide an analysis using established neuroscientific and brain-computer interfacing methods of a sample data set created with the toolbox.

## 2 SEREEGA Architecture and Functionality

### 2.1 Principles of EEG Simulation

To simulate EEG data, the toolbox solves the forward problem of EEG, prescribing how activation signals from specific sources in the brain are projected onto an array of electrodes on the scalp. In matrix notation, this can be written as

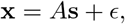

with x denoting the vector of the recorded or simulated scalp signal, s the source activation signal, *A* the projection matrix used to project signals from the source to the scalp electrodes, and *ϵ* denoting a vector of noise.

In SEREEGA, the user defines s, *A*, and *∊*, allowing EEG data x to be simulated.

**Table 1:** Notation used in Section 2.1.

| Notation |  | Description |
| --- | --- | --- |
| $\hat{A} := \mathbf{A}_{\cdot, h, \cdot}$ | $\in \mathbb{R}^{n_{chs} \times 3}$ | Projection matrix of one source |
| $\mathbf{A}$ | $\in \mathbb{R}^{n_{chs} \times n_{srcs} \times 3}$ | Projection tensor of all sources |
| $e \in \{1, \dots, n_{eps}\}$ | $\in \mathbb{N}$ | One of $n_{eps}$ epochs |
| $\mathbf{E}$ | $\in \mathbb{R}^{n_{chs} \times n_{tps} \times n_{eps}}$ | Simulated sensor noise |
| $\mathbf{g}_c^k$ | $\in \mathbb{R}^{n_{tps}}$ | Activation signal of one signal class $c$ for one component $k$, $c \in \{1, \dots, n_{cls}^k\}$ |
| $h \in \{1, \dots, n_{srcs}\}$ | $\in \mathbb{N}$ | One of $n_{srcs}$ sources in the lead field |
| $k \in \{1, \dots, n_{cmps}\}$ | $\in \mathbb{N}$ | One of $n_{cmps}$ brain components |
| $n_{chs}$ | $\in \mathbb{N}$ | Number of channels |
| $n_{cls}^k$ | $\in \mathbb{N}$ | Number of classes for component $k$ |
| $\mathbf{o}^h$ | $\in \mathbb{R}^3$ | Orientation vector for source $h$ |
| $\mathbf{s}^k$ | $\in \mathbb{R}^{n_{tps}}$ | Activation vector of component $k$ |
| $\mathbf{s}_t^k$ | $\in \mathbb{R}$ | Activation of component $k$ at time point $t$ |
| $\hat{\mathbf{s}}_t^k$ | $\in \mathbb{R}^3$ | Oriented activation vector of $\mathbf{s}_t^k$ |
| $t \in \{1, \dots, n_{tps}\}$ | $\in \mathbb{N}$ | One of $n_{tps}$ time points per epoch |
| $\hat{\mathbf{x}}_t^k$ | $\in \mathbb{R}^{n_{chs}}$ | Projected activation of component $k$ at time point $t$ |
| $\hat{X}^k$ | $\in \mathbb{R}^{n_{chs} \times n_{tps}}$ | Matrix of projected activation of component $k$ |
| $X^e$ | $\in \mathbb{R}^{n_{chs} \times n_{tps}}$ | Matrix of projected activations of one epoch $e$ |
| $\mathbf{X}$ | $\in \mathbb{R}^{n_{chs} \times n_{tps} \times n_{eps}}$ | The final simulated EEG data at sensor level |

Source activations are defined on the basis of so-called *components:* for each component, any number of different signal *classes* can be defined, which prescribe how corresponding activation signals are to be generated. For each signal class, a type of activation signal (for example, an event-related potential; see Section 2.3.3) is defined, as well as all corresponding parameters. Furthermore, for each component, a source from the lead field is specified, which is modelled as a dipole at a specific location in the brain. The component also contains this dipole’s (i.e. this source’s) orientation. A component thus prescribes what signal is to be simulated, and how it is to be projected onto the scalp.

SEREEGA simulates one segment of EEG data, or one *epoch*, at a time, and repeats this *n*_*eps*_ times to obtain a larger data set. In the following, we first consider a single epoch. The activation signal of each class, 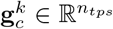, is considered as a vector which consists of the activation signal (i.e. the amplitude time series) for all *n*_*tps*_ time points in this epoch.

Thus, for each component *k*, its activation *s*^*k*^ consists of the summed activations of all signal classes 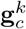 assigned to that component, which are 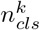 many:

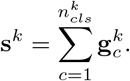

For most signal classes, its exact activation signal 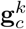 is determined procedurally at runtime based on the specified parameters.

We now describe how an activation signal is projected to the channels. For this, we consider the signal at a single time point *t*, by taking the *t*-th entry of *s*^*k*^, denoted by 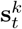. We project this signal from a source denoted by *h*.

The activation signal is projected onto the electrodes on the scalp through a projection matrix *Â* which we obtain using lead field theory (Ferree & Clay, 2000). A lead field contains projection parameters that indicate for each electrode and source how an activation at that source is scaled when it is recorded at that electrode on the scalp. To that end, the activation signal is split in three directions (represented by the three base vectors of Euclidean space), resulting in a third-order tensor consisting of *n*_*chs*_ layers, *n*_*srcs*_ rows and 3 columns:

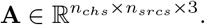

Each row describes the projection matrix for a single source. For a specific source *h* corresponding to a component *k*, 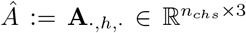 thus describes the projection matrix of that component. The orientation **o**^*h*^ of source *h* can be expressed as a linear combination of the different base vectors contained in the three columns of *Â*:

(1, 0, 0)^*T*^: the x-direction, pointing to the left ear,
(0, 1, 0)^*T*^: the y-direction, pointing to the nose, and
(0, 0, 1)^*T*^: the z-direction, pointing to the top of the head.

This can be expressed by

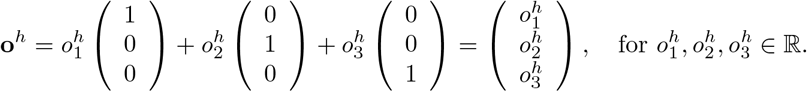

To project a component’s activation onto the scalp, this activation 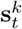 is first oriented by scaling it in the corresponding directions, yielding the oriented activation vector 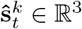:

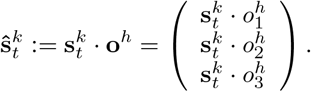

Then, the oriented activation vector 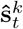 is projected through the lead field which corresponds to the source of the activation, by multiplication with the projection matrix 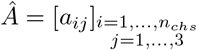:

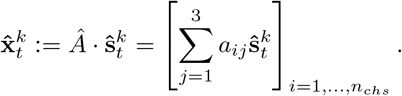

This yields a vector 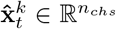 in which every element corresponds to the simulated signal amplitude at time point *t*, projected from source *h* to one electrode. To obtain the corresponding matrix for all time points for the component *k*, the vectors of all time points are concatenated:

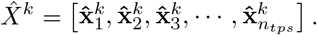

All simulated and projected activation signals 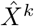 of each component are summed to form one epoch. Thus, for one epoch, the simulated scalp signal *X*^*e*^ has the following form:

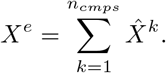

The parameters in the signal classes describe how each component’s activation signals vary between epochs. The projection matrix *Â* can also be made to vary from epoch to epoch, to simulate non-stationarities in the components’ projections, for example due to shifts in electrode positions. Finally, all scalp signal matrices for all epochs are concatenated in the third dimension, yielding a third-order tensor in 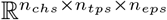. It is at this point that sensor noise **E** is optionally added (source-level noise is defined as an element of 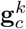). This yields the tensor of simulated EEG data:

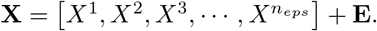

This is the data format used by most software packages to represent epoched EEG data.

**Figure 1:**
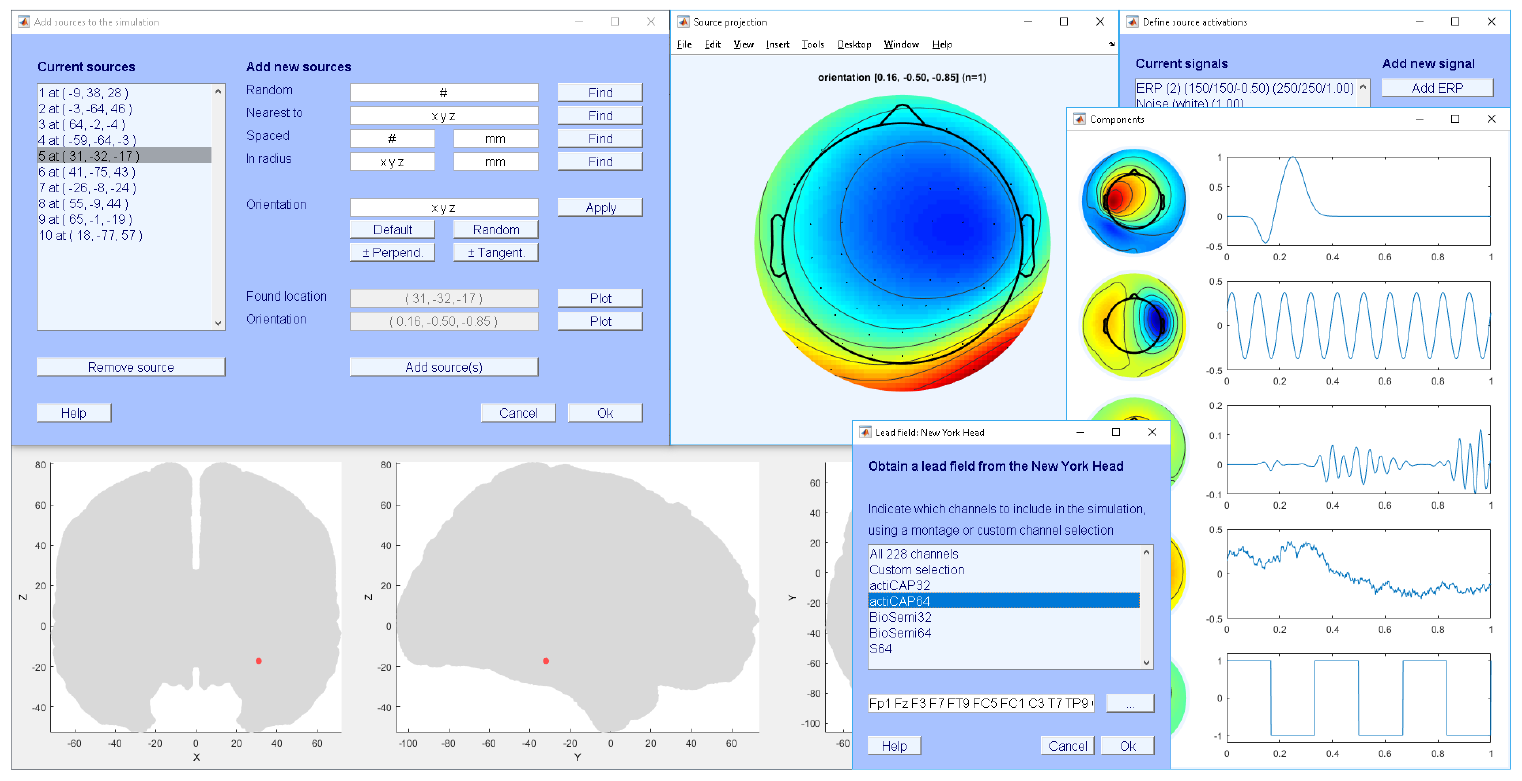
Impression of the SEREEGA GUI.

### 2.2 Platform and License

MATLAB R2014b or higher is recommended for SEREEGA. Some optional functions depend on the Digital Signal Processing (DSP) toolbox version 8.6 (R2014a) or higher. EEGLAB (Delorme & Makeig, 2004) is required as it is used for a number of functions. Lead field generation either requires additional head model files which can be downloaded from their respective websites, or the FieldTrip toolbox (Oostenveld, Fries, Maris, & Schoffelen, 2011). Since SEREEGA is modular, future functions may have further dependencies.

SEREEGA is licensed under the GNU General Public License, version 3, and the code is publicly available on GitHub^1^.

### 2.3 Terminology and Workflow

SEREEGA is available as an EEGLAB plug-in including a graphical user interface (GUI) that allows the core steps of designing and running a simulation to be performed; see figure 1. For more advanced use, SEREEGA is based on written commands and assignments.

A general configuration variable holds information about the number of epochs to simulate, their length, and their sampling rate. This configuration, as well as many other variables, are contained within MATLAB structure arrays.

SEREEGA’s forward model is contained in a *lead field* structure array. The lead field structure contains all possible sources within a virtual brain, modelled as dipoles at specific locations along with their projection patterns. The word *source* thus corresponds to an index in this lead field. For each such index, the lead field contains a position (x, y, z), and projection patterns along three axes (x, y, z). As mentioned in Section 2.1, a linear combination of the three projection patterns can effectively be used to virtually define the source dipole’s *orientation* in 3D space. A source together with its orientation can be saved as a *component* structure array, which additionally needs to be assigned a *signal*: the simulated neuro-electrical activity that will be projected from it, such as an event-related potential or an oscillation at a specific frequency. These are defined separately and added to the components.

The main workflow consists of defining any number of such components, each containing at least one source, orientation, and signal. The simulation of scalp data finally generates each component’s signals and projects them onto the scalp.

The following sections go through the main steps in more detail. Note that these steps must not necessarily be followed in this order.

**Figure 2:**
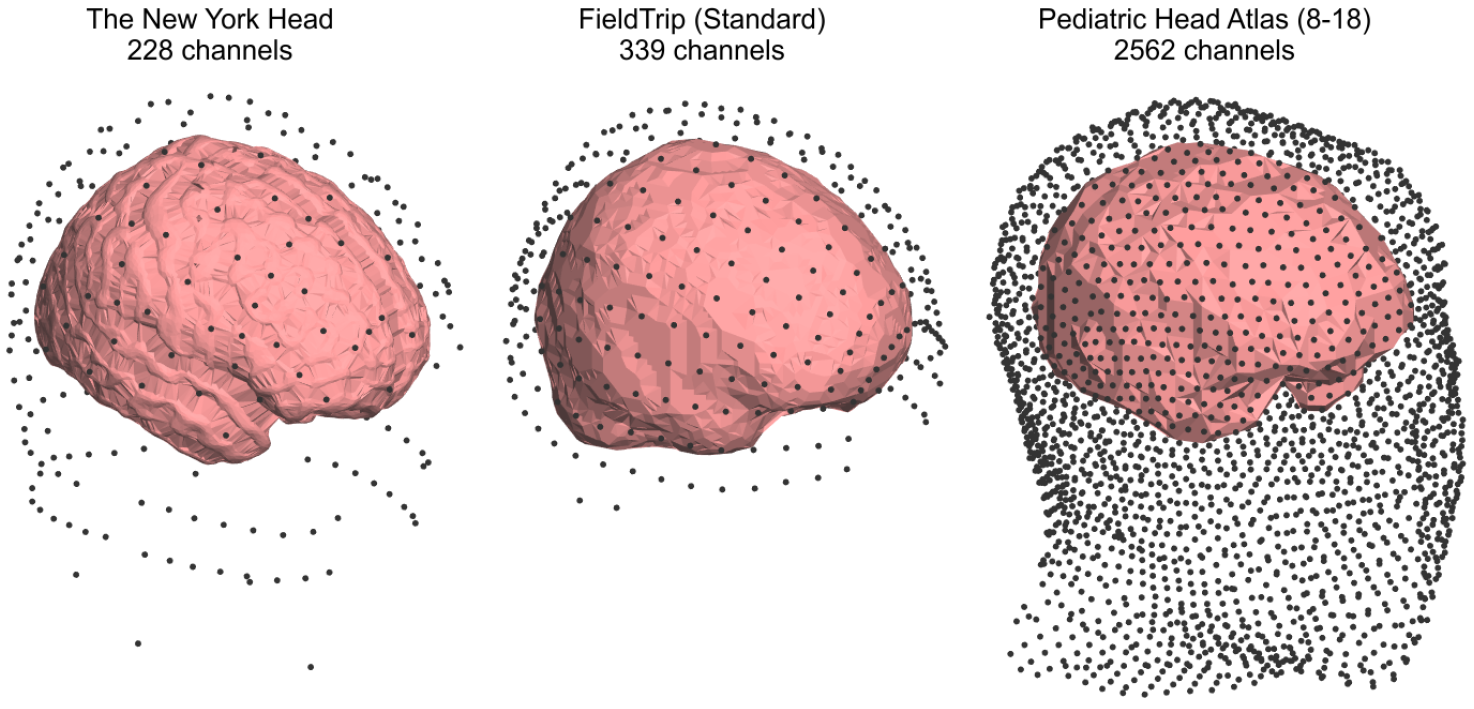
Channel locations and estimated brain boundaries for three head models. Left: The New York Head with its 228 channels. Middle: a 339-channel layout around a standard average head model available by default in FieldTrip. Right: The Pediatric Head Atlas (8 to 18, version 2) showing 2562 channels.

#### 2.3.1 Lead Field Generation

The lead field determines both the maximum possible number of sources as well as the number of channels that will be simulated. Currently, SEREEGA supports two processes to obtain a lead field: it can be obtained from an existing, pre-generated lead field, or a custom lead field can be generated from a given head model. Currently, support for four pre-generated lead fields is included.

The New York Head (ICBM-NY) pre-generated lead field includes almost 75 000 source locations and their projections onto to 228 channels (Huang, Parra, & Haufe, 2016). The electrode positions follow the international 10-05 system (Oostenveld & Praamstra, 2001) but also include two rows of channels below the ears. Each source also contains a default orientation that orients it perpendicular to the cortical surface.

The Pediatric Head Atlases comprise three different head models with pre-generated lead fields for up to three different electrode layouts each, ranging from 128 to more than 2500 electrodes (Song et al., 2013). The models cover heads from three pediatric age clusters, 0-2, 4-8, and 8-18 years old. Lead field sources are spaced in an approximately 1 × 1 × 1 mm grid and range from 3188 to 4837 in number. These lead fields do not contain default dipole orientations. For inclusion in SEREEGA, the given electrode and dipole coordinates are transformed upon initialisation to be centred around (0, 0, 0) and aligned to the axes.

FieldTrip (Oostenveld et al., 2011) can be used to generate a lead field as needed. Using a given head volume, it can generate any number of sources with a given resolution, projecting to any number of channels. By default, a standard head model and a 339-channel definition file following the international 10-05 system are included. FieldTrip does not provide default source orientations.

When obtaining a lead field, it is possible to only use a subset of the available channels by indicating their labels. Convenience functions are available to simulate standard EEG cap layouts with e.g. 64 channels. Figure 2 shows the available channel locations for three head models.

#### 2.3.2 Source Selection and Orientation

The lead field structure contains all possible sources. From these, one or more sources can be selected using different methods to be included in the simulation. It is possible to simply obtain a random source, or to obtain multiple random sources that are at least a given distance apart from each other. It is also possible to select the source nearest to a specific location in the brain, or all sources within a certain radius from a specific location or another source. Sources are referenced using their index in the lead field. Figure 3, left, shows the location of one source using EEGLAB’s standard head model and corresponding plotting function. (In this case, EEGLAB’s standard head model fits the lead field’s head model. SEREEGA’s default plotting functions do not depend on this being the case.)

**Figure 3:**
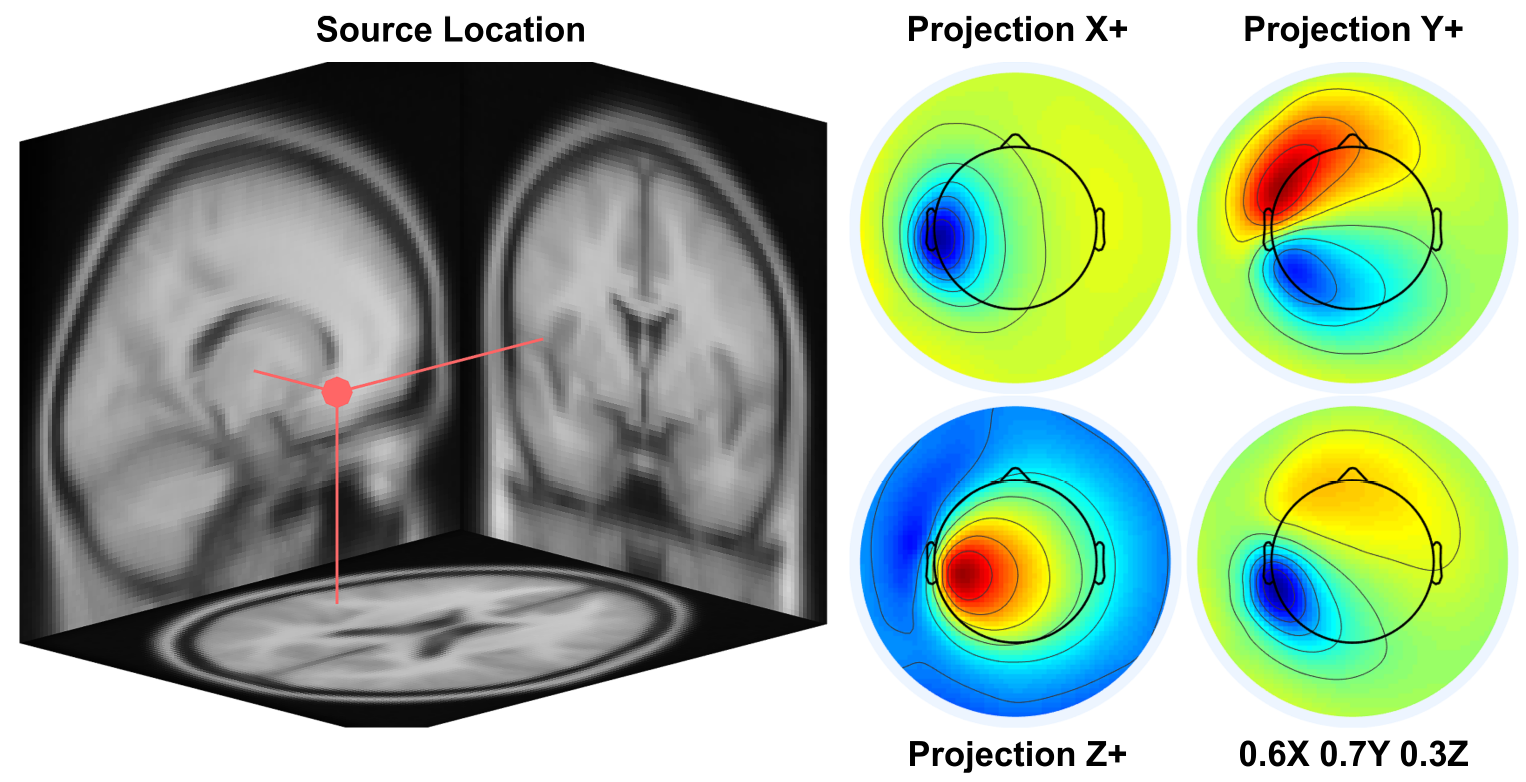
Left: The location of source number 7479 in the New York Head as plotted onto EEGLAB’s standard head model. Right: Four projection patterns of that source onto the scalp. From top left to bottom right: projection in the x-positive, y-positive, and z-positive direction, and a linear combination of the three (arbitrary units). This effectively orients the dipole at the source’s location in 3D space.

Sources represent dipoles at the indicated location. Having selected a source location, this dipole can be oriented to face different directions, which determines how it projects onto the scalp. The lead field makes this possible by containing the dipole’s projection pattern onto all selected electrodes into three mutually perpendicular directions x, y, and z—the base vectors mentioned in Section 2.1. A linear combination of these three projection patterns can then be used to effectively orient the dipole in space. Thus, a dipole’s orientation is indicated using three values, with (1,0,0) representing a perfect x-positive orientation, (0,1,0) y-positive, 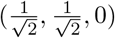 a 45-degree angle with the same amplitude, et cetera. Figure 3, right, shows the three separate patterns as well as a combined projection pattern for one source.

#### 2.3.3 Signal Definition

Having selected a source location and orientation, i.e. a projection pattern, one or more *signals* can be defined to be projected onto the scalp as if they originate from that location. These are defined using *classes*. A class contains a signal’s type and any parameters corresponding to that type in a structure array. For each simulated epoch, a signal will be procedurally generated using the parameters of that class.

SEREEGA currently includes four types of signals: systematic deflections in the time domain (i.e., event-related potentials), systematic modulations of oscillatory activity (i.e., event-related spectral perturbations), stochastic processes in the time domain (i.e., noise), and autoregressive signals (i.e., spatio-temporal signals with systematic interactions between them). Figure 4 provides an overview of three of these and their main parameters. A final *data* type enables the inclusion of any already existing time series as an activation signal. The following sections mention the primary free parameters for each type.

**Event-Related Potentials** An *event-related potential* (ERP; Luck, 2014) class defines one or more positive and/or negative ‘peaks’ or deviations from a baseline in sequence. Each peak is determined by its latency in ms, its width in ms, and its amplitude in μV. For example, the ERP in figure 4 is a single positive peak at 500 ms, 200 ms wide, of 1 μV.

A peak is generated by centring a normal probability density function around the indicated latency with the given width covering 6 standard deviations. This is then scaled to the indicated amplitude. For multiple peaks, each peak is generated individually and then summed together.

**Event-Related Spectral Perturbations** Where ERPs represent systematic activity in the time domain, *event-related spectral perturbation* (ERSP) classes are defined primarily using spectral features. The term ERSP refers to event-related spectral power changes (Makeig, Debener, Onton, & Delorme, 2004); in SEREEGA, it refers to the class of signals that generate specific spectral powers and/or changes therein.

First, a base frequency is defined, either as a pure sine wave of a single frequency, or as a frequency band. In this latter case, uniform white noise is band-pass filtered in the indicated band using a window-based finite impulse response filter with a Kaiser window and an automatically decided filter order (Oppenheim & Schafer, 2010). The left middle panel of figure 4 shows an oscillation of 10 Hz, the panel to the right of that shows filtered noise with power focused between 5 and 10 Hz.

The minimal definition of an ERSP class consists of a frequency (or frequency band) and an amplitude. In case of a single frequency, a phase can optionally be indicated as well.

In other cases, these parameters merely define a base oscillation that is further modulated. The lower row of figure 4 presents two types of modulations that can be defined for ERSP classes: (inverse) burst modulation and amplitude modulation. The former attenuates or amplifies the signal to a given degree using a Tukey window of given width, latency, and tapering. A Tukey window can be adjusted ranging from a square to a cosine shape. This mimics event-related synchronisation and desynchronisation. The latter modulates the amplitude of the base signal to a given degree according to the phase of a sine wave, whose frequency and phase can be defined in the class. Additionally, a pre-stimulus period can be indicated such that the modulation only starts at a given latency.

**Noise** A class of a *noise* signal simply produces coloured noise, from either a normal Gaussian or a uniform distribution, of a given amplitude. A noise class is defined by at least a colour and an amplitude. The upper right panel of figure 4 shows white noise with an amplitude 1 μV.

The Gaussian function uses MATLAB’s DSP toolbox to generate noise with a spectral characteristic of 1/*f*^*n*^, with *n* either −2, −1, 0, 1, or 2. The uniform function first samples the signal from a uniform distribution and then applies a Fourier transformation to achieve the same spectral characteristic.

**Data** For the inclusion of any given time series, for example data generated elsewhere or taken from existing EEG recordings, a *data* class can be defined. It must be given a matrix of data, epoch-dependent indices, and an absolute or relative amplitude. It will project the indicated data scaled to the given amplitude for each simulated epoch.

**Autoregressive model** A final type of class generates signals based on a linear *autoregressive model* (ARM), following the approach by Haufe and Ewald (2016). For a single source activation, this generates a time series where each sample depends linearly on its preceding samples plus a stochastic term. At least the order of the model, i.e. the number of non-zero weights on the preceding samples, and the signal amplitude must be configured. The exact weights of the model are determined automatically.

It is also possible to simulate connectivity between multiple sources. Uni-and bidirectional interactions can be indicated such that a time-delayed influence of one source on another is included in the signal. Either the number of interactions is configured and the toolbox will randomly select the interactions, or a manual configuration of the exact directionalities can be indicated. The model order is the same for all interactions.

Interacting signals are not simulated at runtime, but generated beforehand and included in the simulation as data classes.

**Figure 4:**
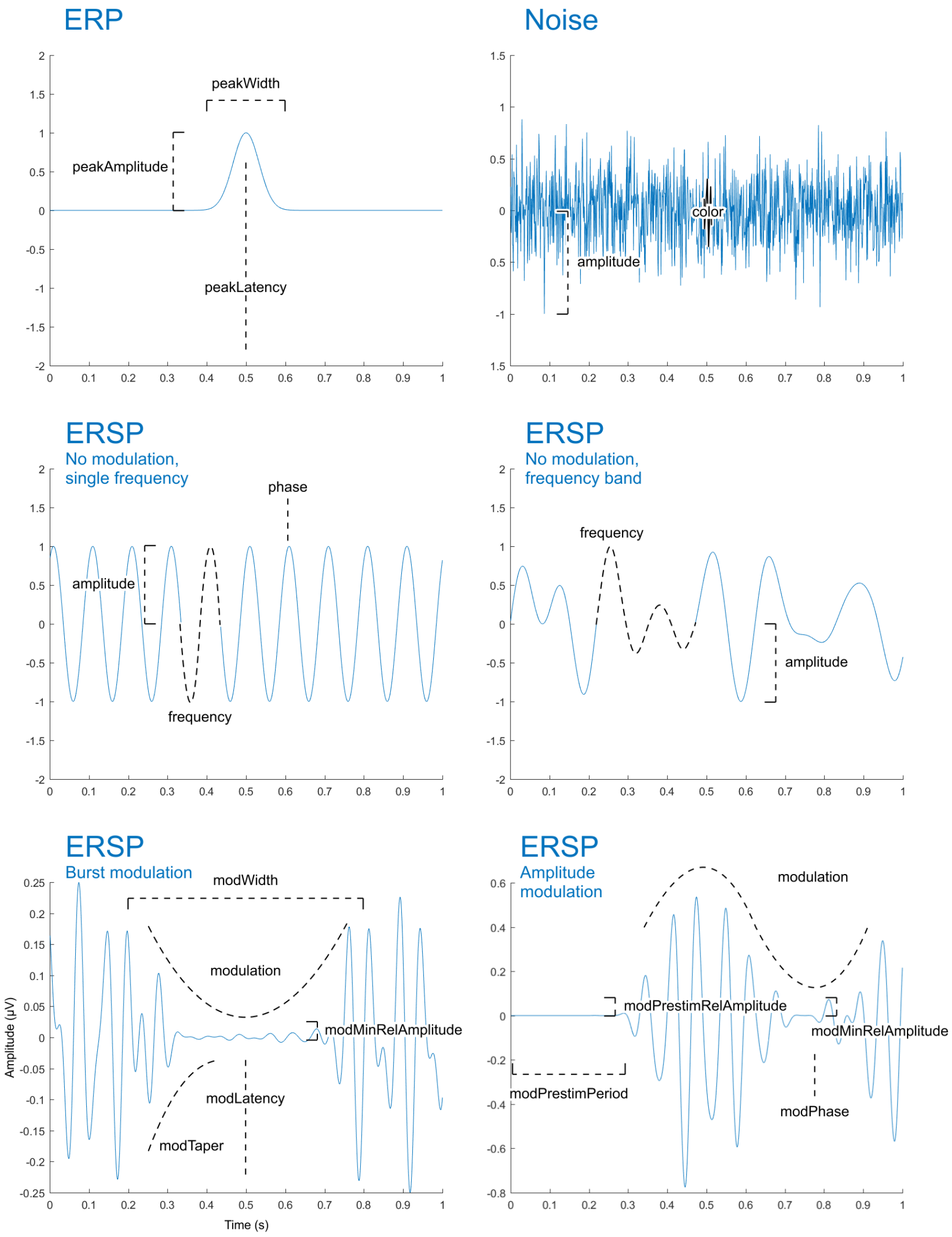
Sample signals annotated with the base variables that control their behaviour. From top left to bottom right: event-related potential (ERP); coloured noise; single-frequency unmodulated event-related spectral perturbation (ERSP); frequency band unmodulated ERSP; inverse burst-modulated ERSP; amplitude-modulated ERSP. Each base variable can additionally be given a deviation and a slope to control the epoch-to-epoch and systematic progressive variability of the signal. Not illustrated: the autoregressive model (ARM) and data classes.

#### 2.3.4 Variability

It defeats the purpose of simulating multiple epochs if every epoch were to be exactly the same. To take an ERP as example, it may not be desirable to have a peak defined at 500 ms appear at exactly 500 ms in every epoch. In fact, perhaps the ERP should sometimes not appear at all. To that end, both classes and components contain parameters to add variability to the generated signal.

For all parameters displayed in figure 4 controlling the shape of the various signals, two more parameters exist that control those base parameters: each parameter has a corresponding *deviation* parameter, and a *slope* parameter.

The deviation indicates a range of allowed values. A base value of 2 with a deviation of 0.3, for example, will vary from epoch to epoch between 1.7 (2 − 0.3) and 2.3 (2 + 0.3) following a Gaussian function. Specifically, the range of deviation covers 6 standard deviations. This can be used to model general Gaussian variability within the system.

The slope indicates a systematic change over time. A base value of 2 with a slope of −0.3, and no deviation, will be 2 on the first epoch, and 1.7 (2 − 0.3) on the final epoch. In between, it will scale linearly with the number of the current epoch. Slopes can be used to model e.g. effects of fatigue and habituation, or to cover a range of intended values.

Each class also has a *probability* parameter that determines, for each epoch, that signal’s probability of appearance. A probability of 0.5 indicates a 50% chance of the defined signal being generated for any simulated epoch. In case it is not generated, a flatline is returned instead, meaning it does not interfere with any other activations at that time.

Deviation and slope parameters are also available for components, to control the dipole orientation. Components can have one additional source of variability, with respect to the source location itself. Instead of one source index, multiple source indices can be indicated as that component’s source. In that case, one of them will be randomly picked for each epoch. As such, together with the orientation variability, spatial variability can be added as well.

#### 2.3.5 Simulating Scalp EEG

A *component* is a structure array that contains, for each element, at least one source index, a corresponding orientation, and at least one signal. Thus, by combining the results of the previous two steps into components, it is specified which of the defined signals are projected from which source at which orientation. When more than one signal is given to one component, these signals are summed together before being projected. It is thus possible to, for example, add noise to an ERP. When the same signal is given to multiple components, a new instance of that signal will be generated for each component at runtime.

A simulation function then takes these components, the lead field, and the general configuration as input, and outputs the simulated scalp data in a *channels* × *samples* × *epochs* matrix, as well as *components* × *samples* × *epochs* source data. For each epoch, this function follows the forward model described in Section 2.1 to produce the scalp data. That is, it generates and sums each component’s signals, projects them onto the scalp using the given lead field, source location, and orientation, and then sums all the projected activations.

At this stage, noise can optionally be added to the final simulated scalp data to simulate sensor noise. This is uniform, temporally and spatially uncorrelated white noise.

#### 2.3.6 Manual and Random Definition of Signal Classes

As a final point it is worth mentioning that not all of the available parameters must necessarily be set by the user. Only a small number are required, and a validation function automatically sets all others to their default values. All deviation-related values default to zero. Non-zero default values are chosen to be most useful; the phase of an oscillation, for example, defaults to a new random value for each epoch. A convenience function exists to add deviations and/or slopes of a certain percentage of each base value to signal classes. When the lead field contains default source orientations, these will be used automatically when no other orientation has been explicitly indicated.

When multiple components are to be simulated, it is also not necessary to manually define all signals. For one, the same signal can be added to multiple components. Also, convenience functions exist to generate any number of ‘random’ ERP or ERSP classes, based on a range of allowed base values.

#### 2.3.7 Sample Code

The following lines of code reflect the workflow explained above, generating 100, 1-second epochs of brown noise from 64 sources spaced at least 25 mm apart, projected onto 64 channels.

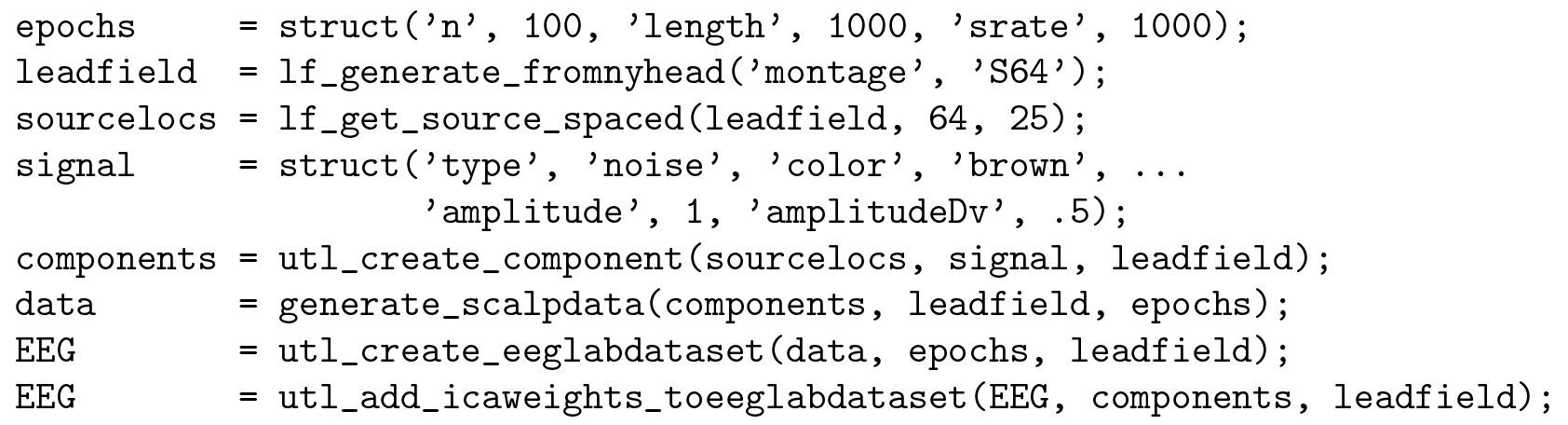

The **`epochs`** structure array contains three fields configuring the number of epochs, their length in ms, and the sampling rate in Hz. On the next line, a lead field is obtained from the New York Head using a preconfigured montage of 64 channels. Next, 64 source locations are randomly selected from the lead field, such that each source is at least 25 mm apart from all the others. As a signal, brown noise is used with an amplitude varying between 0.5 and 1.5 for each epoch. 64 components are then created by combining that same signal with the 64 previously-defined source locations. Because the New York Head is used, no orientations must be indicated explicitly: the toolbox will use the default values included in the lead field. Finally, scalp data is generated: **`generate_scalpdata`** simulates the components’ signals, projects them according to their respective orientations through the lead field, and generates the data set as configured in **`epochs`**. In the last two lines of code, the data set is transformed into an EEGLAB-compatible format, and a ground-truth ICA decomposition is added.

## 3 Sample Data Set

To demonstrate some of the toolbox’s basic functionality, we now present a sample data set that was generated using SEREEGA and simulates a number of known cortical effects during a hypothetical experiment. We then apply established analysis methods to this data and present the results.

### 3.1 Simulated Experiment and Cortical Effects

The hypothetical experiment represents a visual go/no-go experiment. The visual stimulus in question is a pattern change, eliciting a N70-P100-N135 complex over occipital sites (Pratt, 2011). In the ‘go’ condition, the pattern change represents a target stimulus eliciting a strong P3a-P3b complex, most pronounced over the parietal-central-frontal midline (Polich, 2007). Upon perception of the target stimulus, the participants initiate a manual response with their right hand, resulting in mu-and beta band desynchronisation over sites covering the contralateral motor cortex.

Specifically, the visually evoked potential complex was modelled after findings by Gomez Gonzalez, Clark, Fan, Luck, and Hillyard (1994) and Pratt (2011). The N70 was a single dipole projecting centrally from the visual cortex. The P100 and N135 projected bilaterally from Brodmann areas BA 17 and BA 18, respectively. The P3a-P3b was modelled after Polich (2007) and Debener, Makeig, Delorme, and Engel (2005) with dipole locations taken from the latter. The manual response, finally, mimics mu-and beta-band findings of manually responded visual targets by Makeig, Delorme, et al. (2004).

The visually evoked potentials were modelled as single-peak ERP classes centred around 70, 100, and 135 ms, respectively, with widths of 60, 60, and 100 ms, and amplitudes of 7.5, 7.5, and 10 μV, as well as amplitude slopes of −2, −2, and −3 to simulate a level of habituation (Pratt, 2011).

The P3a is an ERP class with a single 400 ms wide peak at 300 ms of amplitude 10 μV, and an amplitude slope of −5, simulating fatigue (Polich, 2007). The P3b ERP consists of two peaks. One at 400 ms (width 600 ms, amplitude 7 μV, amplitude slope −5), and one at 500 ms (width 1000 ms, amplitude 2 μV, amplitude slope −1). This latter peak represents the longer-lasting effects of the P3. The stronger slope of the P3b as compared to the P3a is a known finding reflecting habituation to the stimulus (Polich, 2007). All parameters of the ERP classes were given a deviation of 20% of their value relative to stimulus onset.

The motor cortex is simulated using four components for each class: both left and right motor cortex components are included, each with mu and beta band effects. In the non-target condition all components simply emit resting state mu (here as 8-12 Hz) and beta (19-26 Hz) activity implemented using the ERSP class with no modulation, and an amplitude of 2 μV. In the target condition, the left motor cortex classes are given an inverse burst modulation centred around 650 ms (mu; width 600 ms, taper .5, relative amplitude .5) and 600 ms (beta; width 500 ms, taper .8, relative amplitude .5) respectively. ERSP parameters had a deviation of 10% relative to stimulus onset, with the exception of latency deviation, which was fixed at 100 ms.

To simulate background processes, brown noise was added to each component (as per Freeman, Ahlfors, & Menon, 2009), with an amplitude of 5 μV. Furthermore, brown noise was also projected from a random selection of source locations, such that the total number of components in the data set was 64, all at least 25 mm apart from each other.

A single ‘base’ participant was defined and simulated using the specified component locations and orientations. For 10 subsequent participants, each component’s location was randomly chosen within 20 mm of the original locations. Dipole orientations were randomly varied to be within ±25% of the original value for each axis.

This data set was simulated using the New York Head Model and projected onto the S64 montage of 64 channels. For each condition (target versus non-target), 100 epochs were simulated of 2000 ms each with a 500 ms pre-stimulus period, at a sampling rate of 1000 Hz.

### 3.2 Analyses and Results

The generated data set was subjected to a number of standard analyses to demonstrate their applicability and to illustrate the data itself.

First, figure 5 shows a single-subject ERP image analysis at three midline electrodes of the ‘base participant’. This clearly shows the N70-P100-N135 complex, pronounced most strongly at parietal-occipital sites, as well as the P3, with the P3a most prominent over fronto-central sites and the P3b more pronounced further centrally and parietally. Also visible is the epoch-to-epoch variability of the ERP signals in time and in amplitude, as well as their spatial differences with peaks pronounced differently at different sites. The effects over time can also be seen, with amplitudes diminishing at different rates.

Second, a cluster-based group-level analysis was performed. Using DIPFIT (Oostenveld & Delorme, 2003), equivalent dipole locations were reconstructed from the data set’s ICA decomposition. These locations as well as the independent components’ calculated ERSPs were used for k-means clustering. Due to the variability of the data as well as the clustering algorithm, manual adjustment was still necessary to create one cluster containing all left motor cortex components. The calculated dipole locations are given in figure 6. As was defined, they vary around a given mean location. Figure 7 shows the mean calculated event-related spectral perturbation for this cluster for both the target (left) and the non-target (right) conditions. The target condition shows clear mu and beta effects similar to those found by Makeig, Delorme, et al. (2004, figure 8).

Finally, the data was subjected to an analysis of classifiability for a hypothetical BCI system to be able to detect the elevated P3 response in the target condition compared to the non-target condition, and, separately, the manual response.

**Figure 5:**
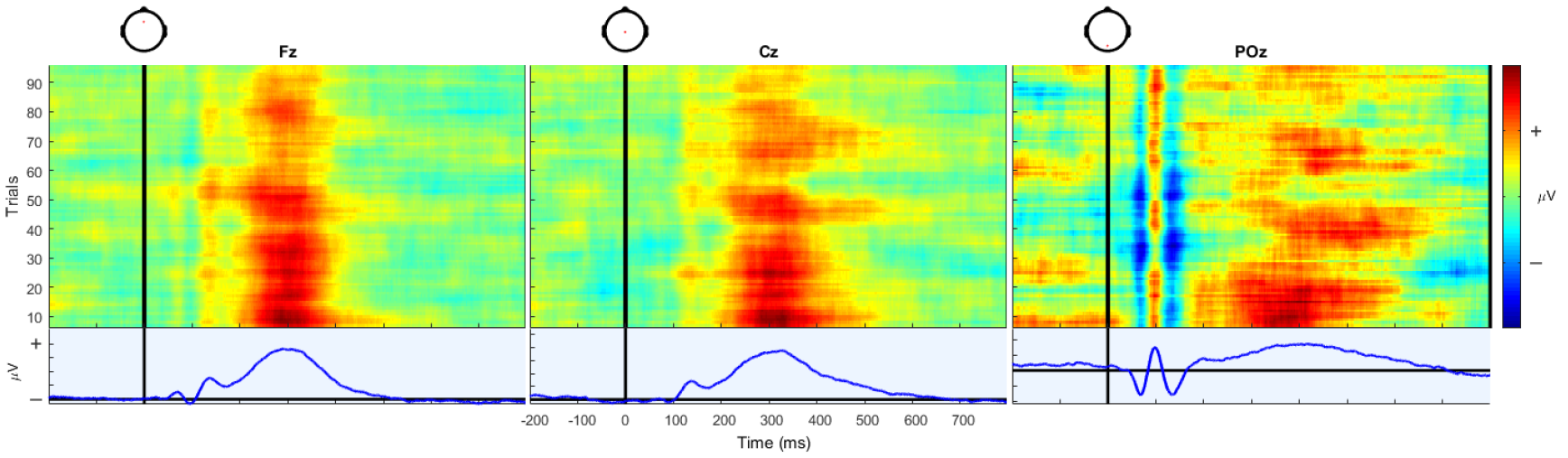
ERP image (Makeig, Debener, et al., 2004) at channel Fz, Cz, and POz showing the epoch-to-epoch variability of the simulated ERP signals as well as their spatial differences and effects over time.

**Figure 6:**
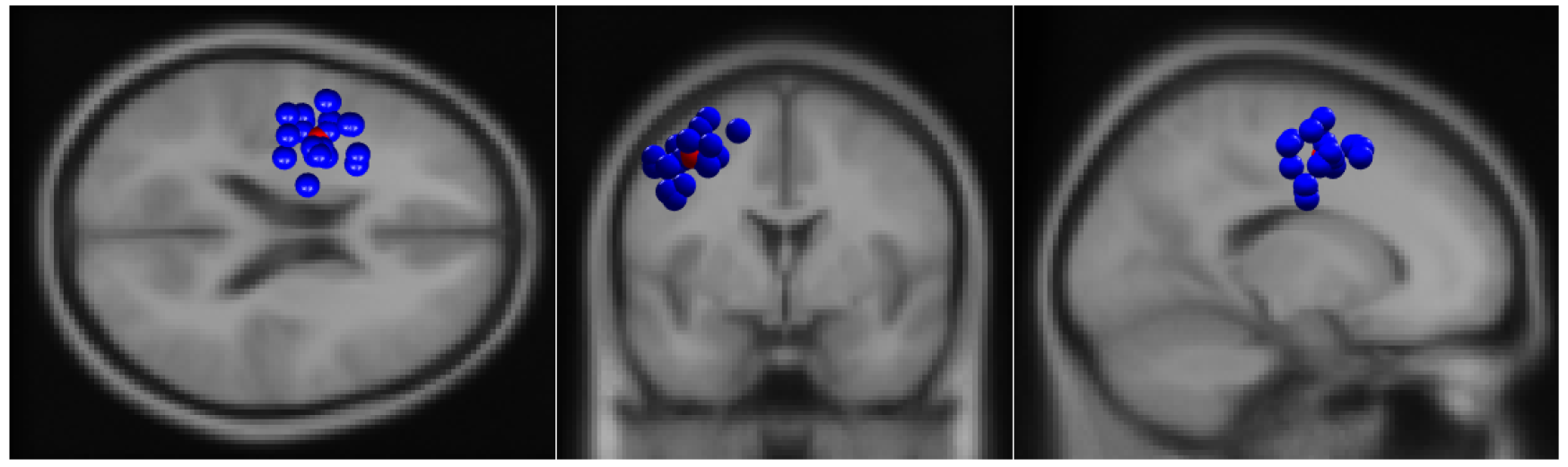
Locations of the left motor cortex equivalent dipoles in each simulated participant, from DIPFIT (Oostenveld & Delorme, 2003) calculations based on the independent components’ scalp maps.

**Figure 7:**
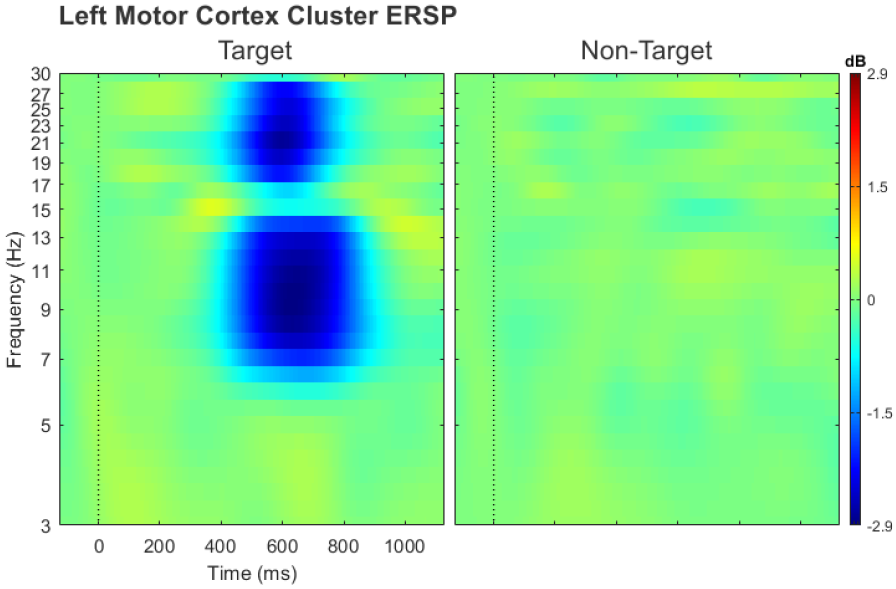
Mean event-related spectral perturbation calculated for the left motor cortex cluster. Left: target condition (right-manual response). Right: non-target condition (no response).

The P3 classifier followed a windowed means approach (Blankertz, Lemm, Treder, Haufe, & Müller, 2011), using the mean amplitude of six consecutive 50 ms time windows starting 200 ms after stimulus onset as features. Before, the signal was bandpass-filtered between 0.1 and 15 Hz.

The manual response was classified using a logBP approach (Pfurtscheller & Neuper, 2001), using as features the power between 7 and 27 Hz, 400 to 1000 ms after stimulus onset, focused on sites covering the motor cortex: FC1, FC2, FC3, FC4, FC5, FC6, C1, C2, C3, C4, C5, and C6.

Both classifiers were implemented using BCILAB (Kothe & Makeig, 2013). A regularised linear discriminant analysis classifier was trained to separate classes (Duda, Hart, & Stork, 2001). A 5 × 5 nested cross-validation with margins of 5 was used to select the shrinkage regularisation parameter, and to generate the estimates of the classifiers’ accuracy listed in table 3.2. With chance level at 50% and significance reached at 57% (Müller-Putz, Scherer, Brunner, Leeb, & Pfurtscheller, 2008), we see that both classifiers reach significance for all ‘participants’, with the exception of participant 5, where no significant difference could be detected between the elicited P3 responses of the two conditions.

This shows the data’s variability on another level and demonstrates that levels of significance can be achieved with neither a ceiling nor a floor effect.

**Table 2:**
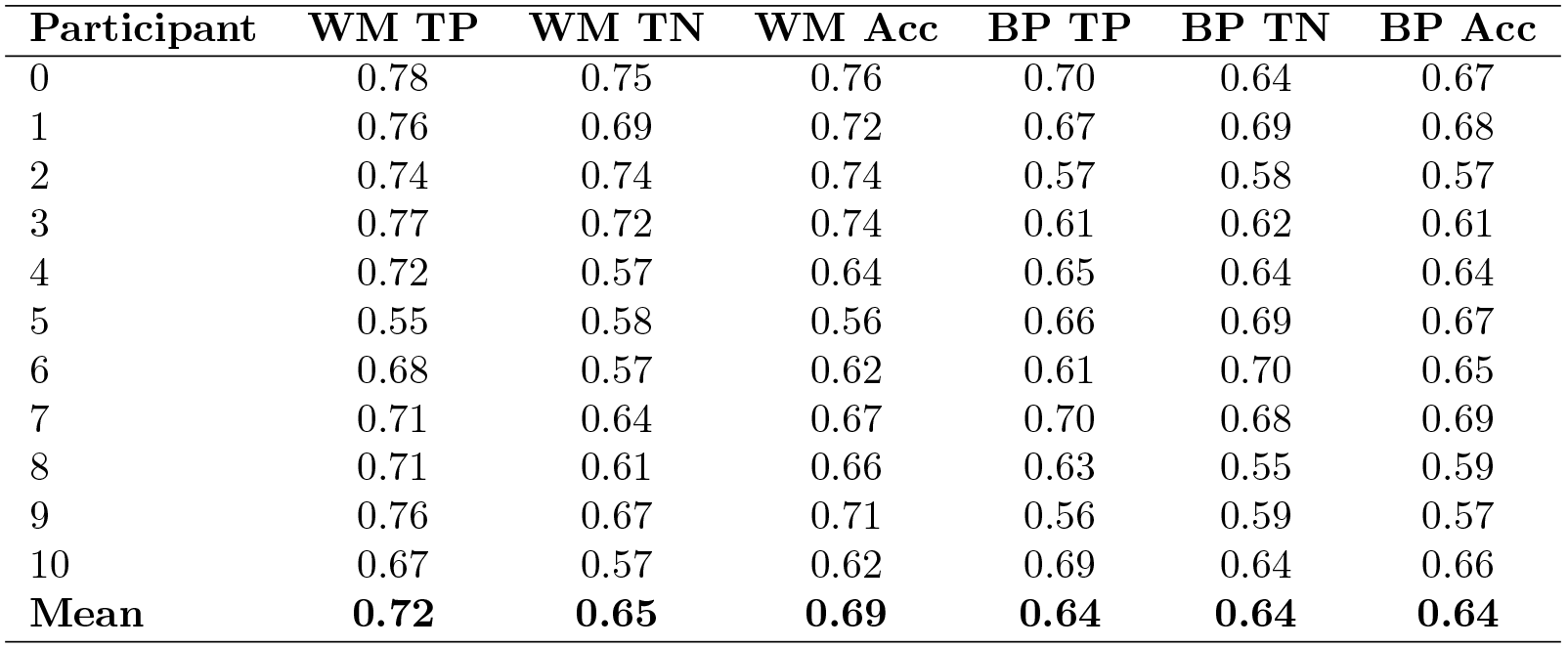
Classification accuracies for two classifiers, one using the windowed means approach (WM) and one based on band power (BP). TP = true positive rate, TN = true negative rate, Acc = overall accuracy. Significance is reached at an accuracy of 0.57. The WM classifier for participant 5 does not reach significance.

| Participant | WM TP | WM TN | WM Acc | BP TP | BP TN | BP Acc |
| --- | --- | --- | --- | --- | --- | --- |
| 0 | 0.78 | 0.75 | 0.76 | 0.70 | 0.64 | 0.67 |
| 1 | 0.76 | 0.69 | 0.72 | 0.67 | 0.69 | 0.68 |
| 2 | 0.74 | 0.74 | 0.74 | 0.57 | 0.58 | 0.57 |
| 3 | 0.77 | 0.72 | 0.74 | 0.61 | 0.62 | 0.61 |
| 4 | 0.72 | 0.57 | 0.64 | 0.65 | 0.64 | 0.64 |
| 5 | 0.55 | 0.58 | 0.56 | 0.66 | 0.69 | 0.67 |
| 6 | 0.68 | 0.57 | 0.62 | 0.61 | 0.70 | 0.65 |
| 7 | 0.71 | 0.64 | 0.67 | 0.70 | 0.68 | 0.69 |
| 8 | 0.71 | 0.61 | 0.66 | 0.63 | 0.55 | 0.59 |
| 9 | 0.76 | 0.67 | 0.71 | 0.56 | 0.59 | 0.57 |
| 10 | 0.67 | 0.57 | 0.62 | 0.69 | 0.64 | 0.66 |
| **Mean** | **0.72** | **0.65** | **0.69** | **0.64** | **0.64** | **0.64** |

## 4 Discussion

In this paper we presented the core functionality of SEREEGA, an open-source MATLAB-based toolbox to simulate event-related EEG activity. SEREEGA allows any number of components to be defined in a virtual brain model, each with a specific location in the brain, a freely oriented scalp projection, and any number of signals. Five different types of activity are currently supported, allowing systematic effects in both the time and frequency domain to be simulated, mimicking known EEG features. Simulated EEG data can serve as ground truth for the development of new EEG analysis methods or the validation of existing ones.

The presented data set serves as a demonstration of some of SEREEGA’s functions: a set of known cortical effects was simulated, mimicking a hypothetical experiment. It furthermore demonstrates how a basic initial configuration can be procedurally varied to simulate any number of additional participants with different anatomical layouts and variations in their cortical responses, leading to different results for example with respect to classification accuracies.

The classification accuracies presented here were only around 7 to 12 percentage points above significance on average. This is primarily due to the relative amplitude of the added noise, i.e., the signal-to-noise ratio of the generated data set. This can easily be varied to increase or decrease the classification accuracies as needed. SEREEGA allows signal and noise components to be simulated separately, and then mixed to achieve a specific signal-to-noise ratio.

Different levels of complexity and realism can be generated using SEREEGA. For some purposes, there may be no explicit need to simulate realistic data sets or otherwise precisely model specific known cortical effects: The results of the methods may hold regardless of the complexity of the simulation. For example, to validate, in theory, a method such as logBP, a pure sine wave among white noise in a low number of randomly mixed channels may suffice. However, other methods do require a higher level of realism with e.g. a higher number of channels and realistic spatial dependecies, a higher number of noise sources, et cetera. For the sake of demonstration, we opted for a more realistic hypothetical experiment to demonstrate in this paper. We do not mean to say however that this level of realism is the explicit goal of SEREEGA, nor that it is methodically preferable. We did intend to illustrates how SEREEGA *can* be used for this purpose. It should also be mentioned that realism here refers to the ultimately simulated source and scalp signals, not their underlying neurophysiology. As such, for example, ERPs are simulated directly as ERP-like time series, not as lower-level phase-resetting oscillations. This level of realism would be unduly complex and not necessary for this toolbox’s purpose.

To the extent that their details were sufficiently reported, all (software) simulations that were performed by the authors mentioned in the introduction of this paper can be done using SEREEGA. The five signal types and other functions of SEREEGA cover and extend the processes used in those methods. This refers to the core steps of each simulation—further implementation details may of course differ. SIFT, for example, being focused on connectivity, has more options to control the parameters of the autoregressive models than SEREEGA currently does.

The separate treatment of signals and components also allows the latter to be flexibly adapted. For example, a spatial Gaussian smoothing of the signal in the head model, such that it no longer comes from a single point but instead from a larger diffuse area in the brain, is easily realised.

Its modularity does add one caveat to the use of SEREEGA. As different head models may use different coordinate formats (e.g. Talairach, MNI, or custom measurements), sources found at the same [*x*, *y*, *z*] location in SEREEGA scripts may correspond to different actual locations depending on the head model used. Although authors who publish head models and lead fields often take great care to standardise their formats, they are not all uniform. Sometimes this is necessary: the dimensions of the Pediatric Head Atlas covering zero-to two-year-olds are of course smaller than those of an adult human’s head model. SEREEGA includes plotting functions to investigate the size, shape, and location of a lead field’s head model and the sources contained within, as well as to compare them against a standard adult head.

Furthermore, the availability of different modules in a general-purpose toolbox places some responsibility with the user to make use of the tools in a way that is compatible with the tested hypothesis. Different analysis methods have different assumptions as to e.g. the number of active sources or the statistical properties of the source signals. While SEREEGA is capable of adhering to a large variety of assumptions, not all data that is produced by it will adhere to the same ones. This depends on the design of the data.

A limitation in the current architecture is that the defined components are necessarily independent at runtime: the procedurally generated activity of one component cannot presently depend on the activity in another. For such interactions to be modelled, the signals must be generated beforehand and included as data classes, as is the case for the autoregressive models.

In its current version, SEREEGA makes five accurate, high-density head models accessible in one toolbox, and allows five types of activation signals, enabling coloured noise, ERP-like deflections in the time domain, oscillatory activity with specific spectral characteristics and/or temporal modulations, and autoregressive signals to be simulated, as well as the inclusion of any other pre-generated time series. This covers and extends the vast majority of past and present-day EEG simulation approaches. Additionally, FieldTrip-generated lead fields based on standard or custom head models can be used, and the toolbox’s architecture allows it to be readily extended with additional head models and signals. For more specific needs, SEREEGA has been designed to be modular with extendability in mind.

SEREEGA thus represents, for the first time, an extensive general-purpose toolbox dedicated to simulating event-related EEG activity, making simulation methods more accessible, standardised, and reproducible.

With SEREEGA, we hope to help developers of EEG-based methods by making data simulation more accessible. With the consistent and increasing popularity of EEG, there is an accompanying need to further develop and validate EEG analysis methods. Simulated data can help with that by providing a ground truth to verify these methods’ results. Seeing the current scope of application of simulated EEG studies, we believe that SEREEGA can help with the great majority of these, and can be easily extended for most of the remaining needs.

## Acknowledgements

Part of this work was supported by the Deutsche Forschungsgemeinschaft (grant number ZA 821/3-1), Idex Bordeaux and LabEX CPU/SysNum, and the European Research Council with the BrainConquest project (grant number ERC-2016-STG-714567). The authors thank Mahta Mousavi for the band-pass filter design, Stefan Haufe for the autoregressive signal generation code and accompanying support, and Ramôn Martínez-Cancino for his assistance with the GUI development.

## Footnotes

1 **https://github.com/lrkrol/SEREEGA**

